# Enhancing Mental Well-being: Efficacy and Neural Mechanisms of Leyi Training in Emotional and Cognitive Intervention

**DOI:** 10.1101/2023.10.10.561370

**Authors:** Xiaofei Jia, Rui Li, Changle Zhou

## Abstract

Leyi mental training comprises physical and mental cultivation methods, such as relaxation, breathing, meditation, koan, and cognitive regulation, with the ultimate goal of achieving enlightenment. Over a decade of practical experience has demonstrated its potential to enhance individuals’ mental well-being. In our study, we employed EEG and psychological scales to investigate the efficacy of Leyi training in emotional and cognitive intervention, as well as its underlying neural mechanisms. The results revealed significant improvements in both emotion and cognition following the intervention. We propose that the effectiveness of Leyi training in emotional regulation may stem from the heightened sensitivity to present experiences and the robust cognitive control ability cultivated through the practice, which enables practitioners to allocate equal, non-reactive attention to all stimuli, regardless of their emotional valence. Moreover, this training facilitates more effective allocation of cognitive resources to deal with current tasks, thus improving cognitive abilities.

## 1. Introduction

Numerous studies have provided evidence of the benefits of various forms of meditation and mindfulness training (Lutz et al., 2007; Walsh and Shapiro, 2006; Tang et al., 2015). From a modern psychological perspective, these methods of mental cultivation significantly contribute to enhancing mental health. The mindfulness method, for instance, is widely employed for stress reduction (Jin, 1992), psychotherapy (Buchheld et al., 2001), addiction withdrawal, among other applications (Vidrine et al., 2009; Chen et al., 2010).

Numerous studies have examined diverse meditation practices and compared their efficacy to non-intervention control groups. The findings indicate that the training cohorts demonstrate superior outcomes compared to the control groups (Jain et al., 2007; Chan and Woollacott, 2007). For instance, Slagter et al. (2007) conducted a study where the experimental group underwent a three-month meditation training program, while the control group was placed on a waiting list and matched the training group in all relevant aspects. The study employed an attentional blink task (Shapiro et al., 1997) to assess participants’ cognitive performance. Prior to the intervention, both groups performed similarly. However, post-meditation, the experimental group demonstrated significant improvement in executive control network performance, indicating the beneficial effects of meditation.

Tang et al. (2015) postulated that meditation operates by enhancing self-regulation, which involves improving attention control, emotional regulation, and self-awareness. This process of self-regulation is in accordance with the concept of “Citta” in Chinese philosophy.

The Sanskrit term “Citta” refers to the practice of nurturing the mind, which is central to Chinese traditional culture and has its roots in the teachings of Confucius. Through rigorous training, individuals can attain an elevated spiritual state (Zhou, 2015). Changle Zhou developed the Leyi Mental Training Method after years of practice, which involves a combination of physical and psychological techniques. Mindfulness meditation is a pivotal component of this approach. The key characteristic of the Leyi Mental Training Method is its focus on achieving the “Tao” as its goal, utilizing Zen and Confucianism as the means of inner cultivation. It merges contemporary psychological adjustments with traditional Chinese Buddhist practices, which include relaxation, breathing, meditation, koan, and disclosure (a form of cognitive adjustment). The ultimate objective is to attain “enlightenment,” a high-level ideology distinguished by emptiness and selflessness (Zhou, 2015).

One of the key stages in Leyi Mental Training is the process of “breaking the ego-mind,” which involves eliminating conceptual distinctions, treating all things equally, and refraining from evaluating situations from the perspective of “self.” Eventually, practitioners can reach a state of non-ego, which is unbound and devoid of self-attachment. Mindfulness meditation is the cornerstone of the Leyi Mental Training Method. It fosters an unreactive sensory perception through attention training and facilitates the transformation of the individual’s consciousness. This shift in consciousness is one of the primary mechanisms responsible for the beneficial outcomes associated with mindfulness meditation (Tang et al., 2015).

In addition to mindfulness meditation, Leyi Mental Training Method also incorporates “case-resolving,” a distinctive element of ancient Chinese Buddhism. During this exercise, the master poses seemingly illogical questions to the disciples, challenging their logical thinking and compelling them to release their attachments. The ultimate aim is to lead the disciples to a heightened state of consciousness known as “zero-reservoir” (which is equivalent to the concept of no-self) (Suzuki et al., 1998). Our diverse training practices, such as meditation and case-resolving, are all intended to achieve this objective.

Recent studies have demonstrated that meditators exhibit increased activation in the frontal regions of the brain (Allen et al., 2012; Lutz et al., 2014). This phenomenon can be attributed to the significant cognitive effort required to attain a meditative state, which involves overcoming habitual reaction patterns. Through intensive training, meditators develop an even-handed approach to emotional and other stimuli, whereby all stimuli are perceived and processed without bias. Consequently, emotional stimuli lose their power to elicit intense emotional reactions.

Mindfulness practice is believed to enhance individuals’ awareness of their present experiences. By genuinely and objectively comprehending their current experience, individuals can more genuinely understand the true nature of things, which is considered to be an enhancement of wisdom (Suzuki et al., 1998; Suzuki, 2017). Through this training, cognitive control in the frontal lobe is enhanced, and attention can be more effectively distributed. This enables individuals to neither indulge nor ignore anything, thereby improving their mental efficiency.

Over a decade of practice has indicated that Leyi training can enhance physical and mental wellbeing (Zhao & Zhou, 2016). However, to present it as scientifically rigorous research, it is necessary to employ quantitative methods and provide valid explanations. Therefore, this study utilized cognitive-behavioral assessment and EEG techniques to investigate the impact of Leyi training and its underlying brain mechanisms.

We enrolled 64 participants who were matched into two equal groups. One group underwent meditation training, while the other served as the control group. Both groups were composed of an equal number of male and female individuals who had no prior experience with meditation. The experimental group received a 7-day full-time on-site training, while the control group was not involved in any training during this period. The training involved various methods, including disclosure, sitting meditation, walking meditation, chanting of mantras, Zen case-solving, and benefit-requesting. These techniques are derived from ancient Chinese Buddhist traditions and are designed to achieve a higher state of consciousness or enlightenment. During the training, the participants followed a strict schedule (see Appendix for the specific daily intervention process), and each training activity had a fixed duration. A group of six individuals practiced intensively with the guidance of an on-site coach.

Both groups underwent a battery of tests one week prior to training and immediately following the final training session. The battery included two parts: a psychological scale test and an EEG test. The present study employed the Profile of Mood States (POMS) to examine the impact of Leyi training on practitioners’ emotional states, while utilizing EEG to investigate the neural mechanisms underlying Leyi training. Specifically, we focused on two EEG measures: the late positive potential (LPP), which is known to be linked with emotion regulation, and the brain network connection, which is associated with cognitive efficiency.

Prior research has posited that meditation enhances self-regulation, which in turn is linked to improved cognitive and emotional regulation (Posner and Rothbart, 2007; Posner et al., 2007). Based on this literature, we hypothesized that: (i) as a result of random allocation, there would be no discernible differences between the experimental and control groups prior to training; (ii) after training, the experimental group would exhibit greater improvements in emotional regulation compared to the control group; and (iii) if cognitive ability were to improve in the experimental group, members would also show higher brain network efficiency, as indexed by small world coefficients (Liang et al., 2010). In simpler terms, we predicted that the experimental group would demonstrate more significant improvements in LPP amplitude, POMS, and brain network organization.

## 2. Methods

### 2.1 Participants

We employed a randomized assignment procedure to assign 32 Chinese participants to an experimental group, with ages ranging from 21 to 40 years and a mean age of 31.9 years, and 32 individuals to a control group, with ages ranging from 19 to 43 years and a mean age of 27.9 years. Both groups comprised an equal number of men and women, and all participants had no prior experience with meditation. Additionally, all participants had normal or corrected-to-normal vision, were right-handed, and reported no history of neurological or psychiatric disorders. Prior to participation, each participant provided written informed consent in accordance with the research protocol approved by the Institutional Review Board of Xiamen University.

### 2.2 Test arrangements

Both the experimental and control groups underwent testing both before and after the training period. All participants took part in an electrophysiological experiment involving EEG recordings, and also completed psychological scales, specifically the Profile of Mood States (POMS).

#### 2.2.1 Electrophysiological experiments

##### 2.2.1.1 Stimuli and Procedure

The stimuli consisted of 126 pictures selected from the Chinese Affective Picture Systems (Bai et al., 2005), with 42 pieces for each emotional category. Valence and arousal ratings for positive, negative, and neutral pictures were respectively (positive pictures: valence=7.01±0.49, arousal=5.19±0.95; neutral pictures: valence=4.95±0.65, arousal=5.01±0.53; negative pictures: valence=2.06±0.67, arousal=5.15±0.83). There was no significant difference in the arousal value of these three types of pictures, but there was a significant difference in the valence.

To present the images, we employed an image perception paradigm, which was based on a previous neuroimaging study conducted by Ochsner and Gross in 2005. The participants were seated in a room that was sound-attenuated and dimly lit, and they were positioned about 0.7 meters away from a computer screen. They were instructed to maintain their gaze on the screen throughout the experiment, with their right index finger placed on the space key.

At the beginning of the experiment, a message prompt was displayed, stating “Please observe the following pictures and assess them immediately upon disappearance from the screen.” The participants initiated the experiment by pressing the space bar. The experiment consisted of three blocks, each of which contained seven units, and each unit was composed of six pictures. The sequence of events for each unit was as follows: a fixation point was initially presented randomly between 800±400ms at the center of the screen, after which a picture was displayed for 1500ms. Participants were then required to indicate whether the image was positive, negative, or neutral by pressing the corresponding F, H, or K key, respectively (F=positive, H=neutral, K=negative). Following their response, the fixation point reappeared, and the second image was displayed, with participants required to respond accordingly. To prevent anticipation of subsequent stimuli, the stimulus interval (ISI) was randomized between 1800ms and 2200ms. Furthermore, within a given unit, images of the same valence were not presented more than two times consecutively. After completion of a unit consisting of six images, a brief rest period was provided before initiating the subsequent unit. Upon completion of seven units, participants were given an extended rest period.

##### 2.2.1.2 EEG recordings

Throughout the recording sessions, the participants were instructed to relax and minimize unnecessary head movements, as well as to avoid blinking during the presentation of stimuli and making responses. EEGs were recorded from the scalp using preconfigured caps that consisted of 64 non-polarizable Ag/AgCl sintered electrodes. The recordings were made using a Neuroscan system, which had a sampling rate of 1000 Hz and a band-pass filter that ranged from 0.1 to 100 Hz. The electrode placement followed the extended 10-20 system. In addition, vertical EOGs were recorded with electrodes placed above and below the left eye. The impedance in all electrodes was less than 5k ohms. The recordings were physically referenced to the nasal tip and were re-referenced offline to the mean of the two mastoids.

##### 2.2.1.3 Data analysis methods

This paper utilized two distinct methods to analyze the EEG data. The first method employed was time-domain analysis, which was utilized to analyze the Late Positive Potential (LPP). The second method involved the use of graph theory to analyze changes in brain networks after training. In the following text, we will introduce and discuss each of these methods in greater detail.

##### Time-domain analysis

EEGLAB 14.1.1 was utilized to analyze the data. Offline, the EEG recordings were re-referenced to the mean of the two mastoids, and each epoch was computed using an 800 ms interval, including a 200 ms period prior to stimulus onset for baseline correction. Band-pass filtering between 0.1 and 30 Hz was applied to the data. Independent component analysis (ICA) was implemented to remove ocular artifacts. The amplitude rejection criterion was set at ±80 µv. Average ERPs were calculated for correct trials that were free of ocular and movement artifacts, with less than 8% of trials rejected due to artifacts. SPSS 17.0 was used for the subsequent statistical analysis, with all p-values obtained from the software output.

##### Brain network analysis

Large-scale brain networks may play a crucial role in supporting the complex state of mindfulness. While traditional EEG signal analysis methods focus on power and amplitude, they do not provide insight into the interactions between different brain regions. As we know, the integration of information across various regions is essential for cognitive processes. Brain network analysis can provide valuable information on the synchronization of EEG signals between pairs of electrodes. By examining changes in brain functional connectivity and topography, the impact of meditation can be quantified.

In 2017, Shaw et al. conducted an investigation into the brain connectivity of meditators and non-meditators utilizing topographic wavelet coherence (Shaw & Routray, 2017). The study revealed a significant difference in topographical connectivity between the two groups, suggesting that meditators exhibit a distinct degree of synchrony among different brain regions during both meditation and the resting state. In 2018, Shaw and Routray compared the efficacy of different measurement methods, such as partial directed coherence (PDC) and directed transfer function (DTF), in assessing EEG signals obtained during meditation (Shaw & Routray, 2018). They observed that, in comparison to DTF, PDC-based brain functional connectivity analysis offers a more comprehensive understanding of the changes in brain topography during Kriya Yoga meditation.

To begin with, EEG data was subjected to a Hamming windowed sinc FIR filter in EEGLAB to produce six-band segments (delta: 0.5-4Hz, theta: 4-8Hz, alpha1: 8-13Hz, alpha2: 13Hz-, beta: -30Hz, gamma: 30-100Hz) in each condition (positive, negative, neutral). The brain networks of each band were then analyzed using a fixed threshold value method. The threshold value was determined based on the EEG data of the pre-training phase, specifically for the positive, negative, and neutral picture conditions. This threshold ensured that the subjects’ brain networks had the same connection density (determined by synchronization likelihood) as the aforementioned states. Using this threshold value, the brain waves of the corresponding states were segmented in the post-training phase to obtain the brain network of these four states.

Next, the small-world coefficients of the experimental and control group’s brain networks in pre-training and post-training were calculated. The difference between the small-world coefficient of post-training and pre-training was computed separately for the experimental and control groups. The difference values of the small-world coefficient were then subjected to ANOVAs to determine whether the changes in small-world coefficients of the two groups were significant.

#### 2.2.2 POMS

Following the EEG experiment, all participants completed the Profile of Mood States (POMS) questionnaire on a computer. Additionally, the control group completed the Life Event Scale. The POMS questionnaire, which comprises 65 items, was administered to both groups before and after the training sessions (Spinella, 2007; McNair et al., 1992). The questionnaire is designed to assess the emotional state of subjects, with each item scored on a five-point scale ranging from 1 (not at all) to 5 (extremely). Seven subscale scores were computed based on the test results, namely tension-anxiety (T), depression-dejection (D), anger-hostility (A), fatigue-inertia (F), confusion-bewilderment (C), vigor-activity (V), and self-related emotions (S), with the first five subscales representing negative mood and the latter two subscales representing positive mood. The internal consistency reliability of the seven subscales is reported to be between 0.85 and 0.87, and the retest reliability is between 0.65 and 0.74 (Liang et al., 1990). Data analysis was performed using SPSS 17.0.

## 3. Results

### 3.1 ERP results

Figure 1 shows the averaged ERP responses using different colors for the six conditions in some representative electrodes. A clear and widespread LPP response was elicited in all six conditions. The Late Positive Potential (LPP) is a salient component frequently analyzed in EEG studies. This positive deflection manifests approximately 300-400 milliseconds after stimulus onset, often localizing to the posterior brain region (Hajcak, MacNamara, & Olvet, 2010; Wessing, Fürniss, Zwitserlood, Dobel, & Junghöfer, 2011). Ample research has demonstrated that changes in the LPP amplitude can signify the degree of emotional arousal during the process of emotional regulation (Wessing, Rehbein, Postert, Fürniss, & Junghöfer, 2013). The distribution of the LPP spans across the frontal, rear frontal, central, parietal, and occipital regions. However, the parietal lobe and central areas have been identified as the most significant, consistent with established LPP features reported in the literature (Hajcak et al., 2010; Wessing et al., 2011). Following previous research, the average amplitude of 12 representative electrodes (F3, Fz, F4, FC3, FCz, FC4, C3, Cz, C4, P4, Pz, P8) was calculated from 300 to 600 ms.

**Figure 1.**
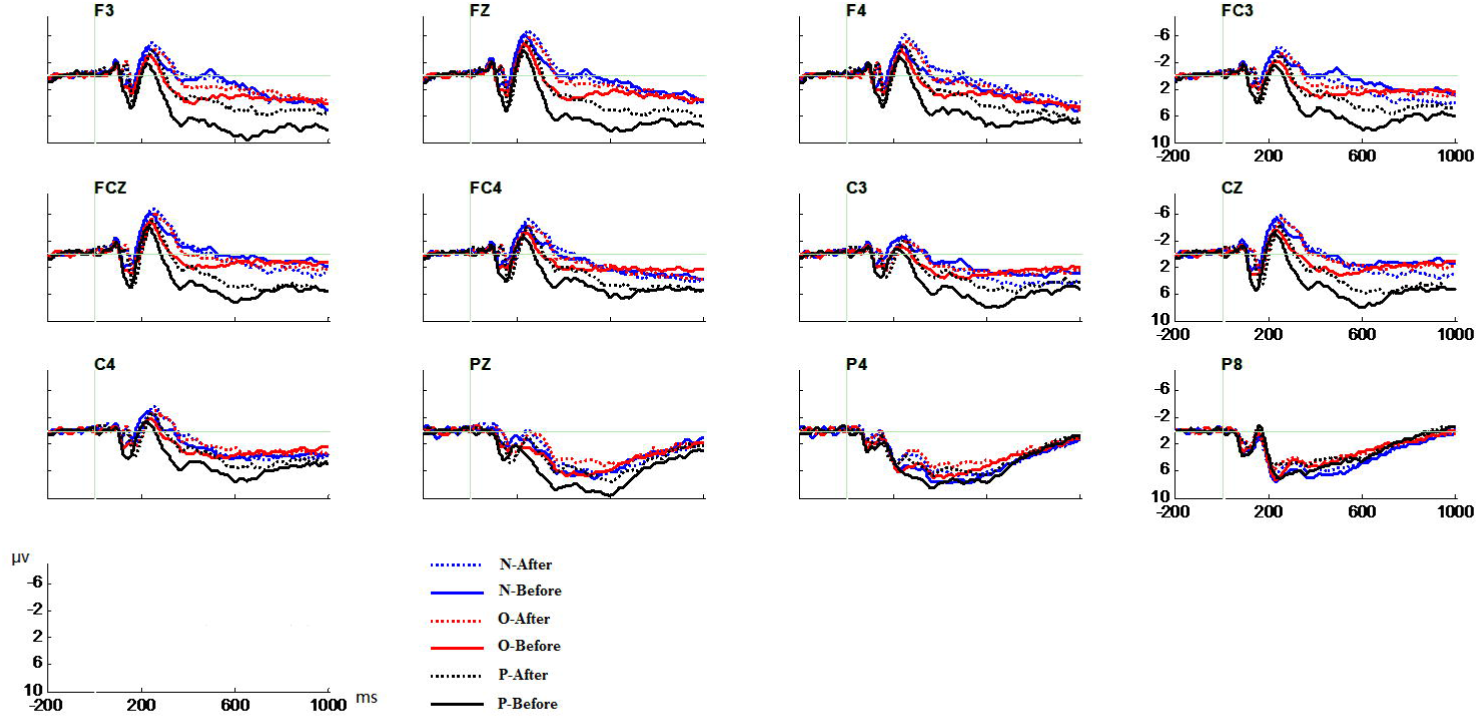
Averaged ERP responses for all conditions are shown in representative electrodes. N-After: the amplitude of negative pictures after training; N-Before: the amplitude of negative images before training; O-After: the amplitude of neutral pictures after training; O-Before: the amplitude of neutral pictures before training; P-After: the amplitude of positive pictures after training; P-Before: the amplitude of positive images before training.

The LPP amplitude induced by positive, neutral, and negative pictures was used as the dependent variable. The group (training and control) and training time (before and after) were used as factors for ANOVAs. Before training, none of the LPP amplitudes of the three types of emotional pictures showed differences between the two groups (P>0.05). The main effect of training is only significant for positive pictures [positive: F (1, 30) = 7.15, P <0.05; neutral: F (1, 30) = 1.52, P >0.1; negative: F (1, 30) = 0.90, P >0.1]. More importantly, the group*time interaction effect is significant for positive pictures [positive: F (1, 30) = 20.79, P <0.001; neutral: F (1, 30) = 1.27, P >0.1; negative: F (1, 30) = 0.42, P >0.1], indicating that the before vs. after differences in the LPP amplitude of positive pictures are significant only for the trained group (see figure 1). The comparison between the pre-training and post-training of the experimental group demonstrated that the LPP amplitude of the three types of emotional pictures all declined. Among them, the positive picture’s decrease is the most significant. The same comparison of the control group didn’t show any significance. This suggests that emotional processing is attenuated following the training intervention, resulting in reduced emotional arousal, particularly for positive emotions (refer to Table 1).

**Table 1.** Comparison of three types of emotional pictures before and after training for the experimental and control groups. *: p<0.05.

### 3.2 Brain network results

First of all, six-band segments (delta (0.5-4Hz), theta (4-8Hz), alpha1 (8-13Hz), alpha2 (13Hz-), beta (-30Hz), gamma (30-100Hz)) were performed on EEG data, then the brain networks of each band were analyzed (see the details in materials and methods section). Thus get the difference value of the small-world coefficient between pre-training and post-training of the experimental and control group. We then performed ANOVAs on these difference values, checking whether the change of the small world coefficient of the two groups was significant. See the results in Table 2 (The p-value of multiple comparisons for six frequency bands has been corrected by Bonferroni).

**Table 2.** Comparison of the difference value of the small-world coefficient between pre-and post-training of the experimental and control group. *: p<0.05.

The results suggest that after training, the brain network of the experimental group became less “small-world,” which indicates a more efficient neural processing. This was observed specifically in the alpha2, beta, and gamma frequency bands, and was not observed in the control group. These findings support the neural efficiency hypothesis, which proposes that training can lead to more efficient neural processing by reducing unnecessary neural activity.

### 3.3 POMS results

The Profile of Mood States (POMS) was employed to assess emotional states in the two groups. Analysis of Variance (ANOVA) was conducted with group (training and control) and training session (pre and post) as factors, using the score of each subscale of POMS as the dependent variable. Prior to training, there was no discernible difference in the six subscales of POMS between the two groups (P>0.05). ANOVA results revealed a group*session effect for subscales A [F (1, 37) = 5.93; P <0.05], D [F (1, 37) = 6.03; P =0.019], T [F (1, 37) = 3.75; P <0.05], and F [F (1, 37) = 5.11; P =0.03]. These four subscales are negative mood scales. It is evident from the results that only these subscales displayed significant differences between the experimental and control groups (see Figure 2). These findings suggest that short-term Leyi training can effectively reduce negative mood states.

**Figure 2.**
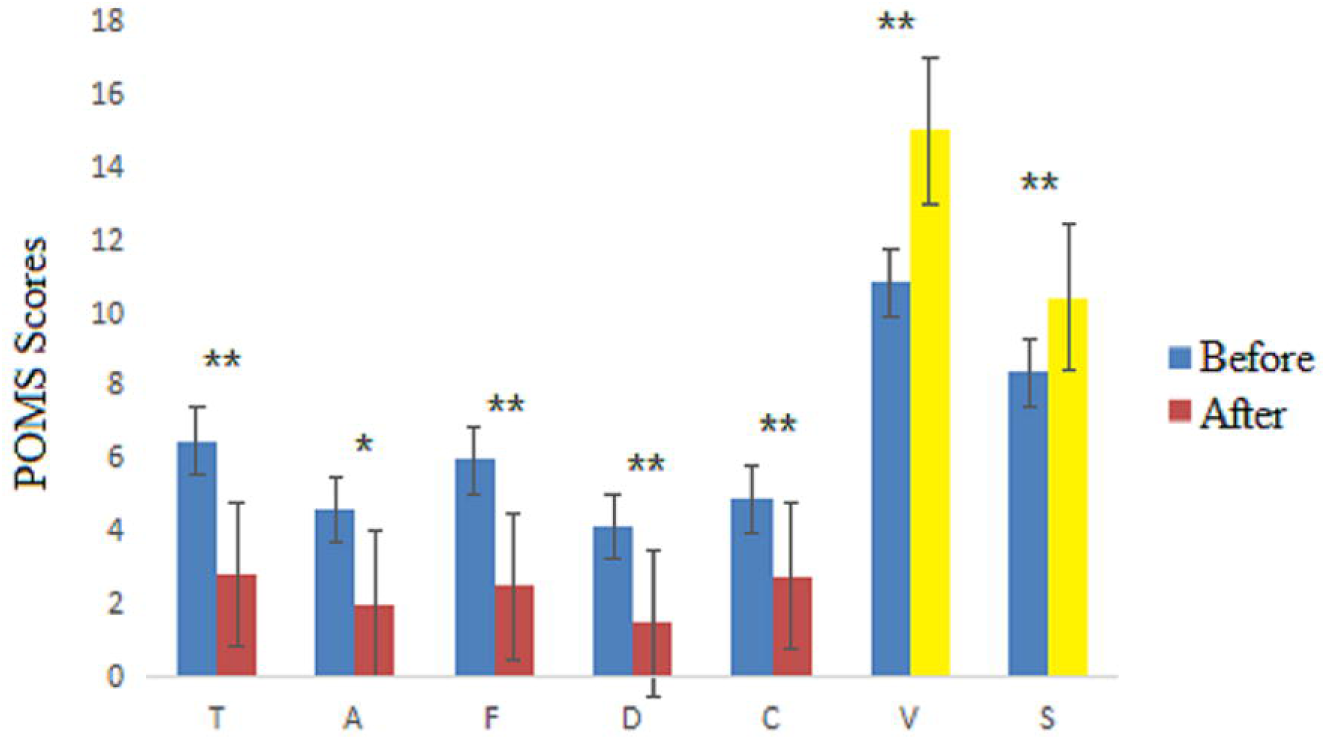
Comparison of the six subscales of POMS before and after training for the experimental group and the control group.

## 4. Discussion

As anticipated, our findings indicate that Leyi training led to more significant improvement in LPP and POMS scores in the experimental group compared to the control group after seven days of training. Since the participants were randomly assigned to either group, we can infer that the training was the cause of the observed emotion regulation improvements. Furthermore, the experimental group demonstrated significant enhancement in cognitive efficiency, as evidenced by a decrease in the small world coefficient relative to the control group. This finding reinforces the potential of Leyi training for cognitive enhancement, consistent with previous studies (Zhao and Zhou, 2016).

The attenuation of the late positive potential (LPP) in response to positive stimuli may be attributed to attentional reallocation following meditation training. Past research has demonstrated that meditation promotes such reallocation, as evidenced by studies on meditators’ attentional blinks. This phenomenon underscores the brain’s limited ability to perceive two successive stimuli in a short span, whereby intense focus on the first stimulus results in missing the second. Slagter et al. (2007) have posited that meditation training can mitigate this fixation on the first stimulus by cultivating non-reactive sensory awareness, thereby reducing the attentional blink phenomenon. As expected, the group of practitioners who underwent high-intensity meditation training for three months demonstrated a higher frequency of detecting two stimuli in quick succession compared to the control group. The decrease in specific brain wave activity in response to the first stimulus among practitioners reflects an enhancement of non-reactive sensory perception. Furthermore, the P3b results indicate that the practitioners can allocate their attention more effectively (Brefczynski et al., 2007; Zeidan et al., 2010; Van et al., 2014).

Other studies have found that the amplitude of the late positive potential (LPP) is associated with valence-driven processing, and LPP selectively reacts only to positive words. Kissler et al. (2013) argue that positive words are related to an enhanced LPP, indicating that positive information attracts more attention and leads to more in-depth stimulus coding than negative and neutral information (Kissler and Herbert, 2009). Herbert et al. (2011) interpreted it as selective processing of positive words, indicating that healthy subjects have a natural tendency towards positive information (Herbert et al., 2008). The results of this study show that meditation training reduces this automated facilitation response. It can redistribute attention, which trains non-reactive sensory perception, paying equal attention to each picture rather than indulging in one of them.

Mindfulness training can improve emotional regulation and reduce stress (as indicated by the results of the Profile of Mood States or POMS), which may also be related to the reaction pattern cultivated in meditation practice - the non-reactive sensory awareness (Malinowski, 2013; Moore et al., 2012). The Leyi training program includes mindfulness meditation, which is known as “open monitor” meditation. The practitioners are always aware of what is happening around them but do not immerse themselves in any particular feeling or idea. Such training can develop non-reactive sensory awareness (Faber et al., 2015; Travis and Shear, 2010; Lutz et al., 2008). In this open meditation process, the brain is in a calm and relaxed state, but thinking remains clear and keen. The practitioners are more acute in their perception of the present, but they do not become attached to any specific things. They can feel pain, but they are not going to explain, change, reject or ignore this feeling (Travis and Shear, 2010). The researchers found that pain did not diminish in the meditators, but the distress caused was not as evident as in the control group.

This finding is consistent with the principles of the emotional ABC theory. According to this theory, the event (A) that triggers an emotion is not the direct cause of the emotional outcome (C). Instead, it is the individual’s belief system (B) and their interpretation of the event that leads to the emotional response. Ellis, the theory’s founder, argued that irrational beliefs cause emotional distress. However, meditation practitioners develop non-reactive sensory awareness, allowing emotional events to occur naturally without judgment or evaluation. As a result, emotional distress is minimized or avoided altogether (Gillani and Smith, 2001; Smith, 2004). This may explain why the experimental group experienced reduced emotional distress after the training.

After seven days of practice, our research has demonstrated the potential of Leyi in regulating emotions, as evidenced by the results of the Profile of Mood States (POMS). The various components of Leyi, such as controlled breathing, body relaxation, and meditation, may be responsible for this effect. Compared to traditional regulation methods that require months of training, Leyi’s simple yet effective techniques can benefit individuals in today’s fast-paced world. Given the high levels of stress that many people experience, even without any diagnosed medical conditions, Leyi’s ability to facilitate physical and mental adjustments is especially relevant.

Large-scale brain networks may underlie the complex mindfulness state. Although researchers commonly analyze EEG signals using power and amplitude, this method does not provide information about the connectivity between different brain regions. Integration of information across various regions is critical in cognitive processing. Brain network analysis can provide information on the synchronization of EEG signals between two electrodes, enabling the study of coupling between different brain regions.

Several studies have found a significant negative correlation between connectivity and IQ in the frontal area (Barry et al., 2002; Martín et al., 2001; Silberstein et al., 2003). High IQ individuals have weaker coupling between different regions, consistent with the neural efficiency hypothesis. This hypothesis proposes that intelligence reflects the efficiency of the brain in recruiting regions related to tasks, rather than irrelevant regions.

EEG studies using various cognitive tasks consistently report a negative correlation between brain activity and intelligence (Barry et al., 2002; Jaušovec, 2000; Riva et al., 2007; Jaušovec N and Jaušovec K, 2003). This finding suggests that the brains of high IQ individuals are less active than those of individuals with average IQ when solving problems. Psychologists have attempted to explain this phenomenon using the Neural Efficacy Hypothesis, which proposes that high IQ individuals typically use brain regions related to the task and avoid irrelevant regions. This hypothesis receives support from research using other methods, such as PET and fMRI (Bechtereva et al., 2004; Rypma et al., 2005; Lee et al., 2006; Micheloyannis et al., 2006; Haier et al., 2003).

Brain network analysis revealed a significant weakening in the small-world coefficient of the experimental group in the alpha and beta bands compared to the control group. This finding supports the neural efficiency hypothesis, indicating that the nervous system’s operating efficiency significantly improves after short-term training. Complex cognitive tasks require a more optimized network structure to complete. After training, fewer mental resources are required.

Additionally, the small-world coefficient of the gamma band in the experimental group showed a significant decline compared to that of the control group after training. The gamma band is a high-frequency band associated with various cognitive processes, such as memory matching and decision-making (Herrmann et al., 2004). Moreover, interconnecting different functional regions in advanced mental processes requires synchronized gamma oscillations (Başar et al., 1996). For example, studies show that learning ability requires gamma oscillations (Miltner et al., 1999). Bassett et al. (2006) studied the topological properties of brain networks (1.1-75 Hz) using ERF technology under various tasks. They found that the brain network under different frequency bands met the “small-world” properties, suggesting that the brain network has self-similar fractal characteristics under those conditions. The study also found that the global topological properties of all bands are relatively stable and not affected by motion tasks. However, in the high-frequency band, the researchers found the emergence of new connectivity and core nodes in the frontal and parietal lobes. This finding suggests that, while maintaining the “small-world” structure, brain networks also spontaneously regulate local connections to meet different requirements. This study found that the small-world characteristics of the brain network weakened in the high-frequency band after Leyi training, indicating that training affected the connection to better adapt to the current information processing. As the participants’ cognitive ability improved after training, weaker neural oscillatory activity was sufficient to complete the same mental tasks.

The present study has demonstrated the potential of Leyi training in emotional regulation and cognitive enhancement. The ultimate objective of Leyi training is to attain a higher level of consciousness characterized by selflessness or emptiness. Therefore, practitioners must invest considerable cognitive effort to overcome their habitual reactive response patterns during the initial stages of practice. This process fosters robust cognitive control abilities, which allow practitioners to treat all experiences in a non-reactive manner, irrespective of their emotional valence, thus demonstrating the effectiveness of Leyi training in emotional regulation. The heightened sensitivity to present experiences developed through this practice enables individuals to perceive things in a more realistic and objective manner, representing a form of wisdom espoused by Buddhist philosophy. Furthermore, our research has corroborated the positive impact of Leyi training on cognitive abilities.

Previous research has highlighted executive attention as a primary mechanism for regulating both cognitive and emotional processes (Posner and Rothbart, 2007; Tang et al., 2015). Studies have suggested that the anterior cingulate gyrus plays a crucial role in executive control (Posner and Rothbart, 2007; Tang et al., 2015; Tang et al., 2007). We anticipate that future investigations will reveal the relationship between Leyi training and alterations in this neural network, elucidating the brain mechanisms underlying the beneficial effects of Leyi training on emotion and cognition.

## 5. Conclusion

Leyi training is a comprehensive approach that incorporates various techniques such as relaxation, breathing, meditation, koan, and cognitive regulation to promote physical and mental well-being. It blends modern psychological adjustment techniques with traditional Chinese Buddhist practices and aims to attain a higher level of consciousness. Our research on the psychological and neural effects of Leyi training led to the following conclusions: 1. The LPP results revealed that Leyi training reduces automatic facilitation responses to positive stimuli and facilitates attentional allocation. 2. POMS results indicated that Leyi training can alleviate negative moods. 3. Brain network analysis demonstrated that Leyi training enhances cognitive efficiency. Overall, our findings support our hypothesis that Leyi training is an effective and accessible method for promoting self-regulation in both cognition and emotion.

## Supporting information

zip

