## Supplementary material for "Enhancing Mental Well-being: Efficacy and Neural Mechanisms of Leyi Training in Emotional and Cognitive Intervention": zip: Table1.docx

**Table 1.** Comparison of three types of emotional pictures before and after training for the experimental group and control group. *:p<0.05

| **Average amplitude** | **Experimental group** | | | | **Control group** | | | |
| --- | --- | --- | --- | --- | --- | --- | --- | --- |
|  | **Before** | **After** | **t** | **Significance** | **Before** | **After** | **t** | **Significance** |
| **Positive pictures(P)** | 4.790 | 2.730 | 4.080 | 0.000* | 5.920 | 4.680 | 1.440 | 0.160 |
| **Neutral pictures(O)** | 2.660 | 1.590 | 1.840 | 0.077 | 2.540 | 2.180 | 0.410 | 0.687 |
| **Negative pictures(N)** | 1.690 | 1.590 | 0.100 | 0.924 | 2.370 | 2.820 | -0.360 | 0.722 |
