## Supplementary material for "Enhancing Mental Well-being: Efficacy and Neural Mechanisms of Leyi Training in Emotional and Cognitive Intervention": zip: Table2.docx

**Table 2.** Comparison of the difference value of the small-world coefficient between pre-training and post-training of the experimental and control group. *: p<0.05

|  |  |  | **Neutral pictures** | | | | **Positive pictures** | | | | **Negative pictures** | | |
| --- | --- | --- | --- | --- | --- | --- | --- | --- | --- | --- | --- | --- | --- |
|  |  |  | **Mean** | **SD** | **Significance** | | **Mean** | **SD** | | **Significance** | **Mean** | **SD** | **Significance** |
| **delta** | Experimental group | Before | 2.22 | 0.08 | | group:  F = 0.002  p = 0.965  time:  F = 0.897  p = 0.347  group*time:  F = 0.003  p = 0.96 | 2.21 | | 0.07 | group:  F = 0.101  p = 0.752  time:  F = 3.714  p = 0.058  group*time:  F = 0.041  p = 0.840 | 2.24 | 0.08 | group:  F = 0.06  p = 0.807  time:  F = 3.925  p = 0.051  group*time:  F = 0.01  p = 0.976 |
|  |  | After | 2.26 | 0.22 | |  | 2.28 | 0.21 | |  | 2.31 | 0.21 |  |
|  | Control group | Before | 2.23 | 0.07 | |  | 2.21 | 0.07 | |  | 2.23 | 0.06 |  |
|  |  | After | 2.26 | 0.22 | |  | 2.30 | 0.30 | |  | 2.30 | 0.20 |  |
| **theta** | Experimental group | Before | 2.24 | 0.08 | | group:  F = 0.06  p = 0.807  time:  F = 3.925  p = 0.051  group*time:  F = 0.001  p = 0.976 | 2.23 | 0.07 | | group:  F = 1.155  p = 0.286  time:  F = 0.036  p = 0.85  group*time:  F = 1.616  p = 0.207 | 2.22 | 0.08 | group:  F = 1.010  p = 0.318  time:  F = 0.682  p = 0.411  group*time  F = 2.021  p = 0.159 |
|  |  | After | 2.31 | 0.21 | |  | 2.18 | 0.11 | |  | 2.16 | 0.14 |  |
|  | Control group | Before | 2.23 | 0.06 | |  | 2.22 | 0.07 | |  | 2.21 | 0.06 |  |
|  |  | After | 2.30 | 0.20 | |  | 2.26 | 0.25 | |  | 2.23 | 0.20 |  |
| **alpha1** | Experimental group | Before | 2.16 | 0.05 | | group:  F = 0.761  p = 0.386  time:  F = 0.174  p = 0.678  group*time  F = 2.221  p = 0.140 | 2.15 | 0.06 | | group:  F = 3.301  p = 0.073  time:  F = 0.027  p = 0.870  group*time  F = 2.849  p = 0.095 | 2.15 | 0.06 | group:  F = 4.115  p = 0.046  time:  F = 0.054  p = 0.818  group*time:  F = 5.315  p = 0.024 |
|  |  | After | 2.12 | 0.12 | |  | 2.11 | 0.10 | |  | 2.09 | 0.10 |  |
|  | Control group | Before | 2.15 | 0.05 | |  | 2.15 | 0.05 | |  | 2.14 | 0.06 |  |
|  |  | After | 2.17 | 0.16 | |  | 2.20 | 0.20 | |  | 2.20 | 0.18 |  |
| **alpha2** | Experimental group | Before | 2.16 | 0.06 | | group:  F = 1.649  p = 0.203  time:  F = 0.124  p = 0.726  group*time  F = 4.470  p = 0.038 | 2.15 | 0.06 | | group:  F = 3.527  p = 0.064  time:  F = 0.005  p = 0.945  group*time  F = 3.986  p = 0.049 | 2.15 | 0.06 | group:  F = 6.228  p = 0.015  time:  F = 0.185  p = 0.668  group*time:  F = 8.530  p < 0.01 |
|  |  | After | 2.10 | 0.12 | |  | 2.09 | 0.10 | |  | 2.07 | 0.10 |  |
|  | Control group | Before | 2.14 | 0.11 | |  | 2.14 | 0.05 | |  | 2.14 | 0.05 |  |
|  |  | After | 2.18 | 0.16 | |  | 2.20 | 0.20 | |  | 2.20 | 0.17 |  |
| **beta** | Experimental group | Before | 2.18 | 0.06 | | group:  F = 6.555  p = 0.012  time  F = 0.053  p = 0.818  group*time  F = 9.558  p <0.01 | 2.17 | 0.06 | | group:  F = 8.510  p <0.01  time:  F = 0.147  p = 0.703  group*time  F = 8.631  p<0.01 | 2.18 | 0.06 | group:  F = 7.607  p <0.01  time:  F = 0.682  p = 0.411  group*time:  F = 12.272  p < 0.01 |
|  |  | After | 2.10 | 0.15 | |  | 2.08 | 0.15 | |  | 2.07 | 0.15 |  |
|  | Control group | Before | 2.17 | 0.05 | |  | 2.17 | 0.06 | |  | 2.16 | 0.06 |  |
|  |  | After | 2.24 | 0.16 | |  | 2.24 | 0.19 | |  | 2.23 | 0.16 |  |
| **gamma** | Experimental group | Before | 2.13 | 0.06 | | group:  F= 6.13  p = 0.015  time:  F = 1.508  p = 0.223  group*time  F = 7.347  p<0.01 | 2.13 | 0.06 | | group:  F = 5.878  p = 0.018  time:  F = 0.757  p = 0.387  group*time  F = 6.774  p = 0.011 | 2.12 | 0.07 | group:  F = 7.627  p < 0.01  time:  F = 0.762  p = 0.385  group*time:  F = 8.656  p < 0.01 |
|  |  | After | 2.08 | 0.17 | |  | 2.07 | 0.16 | |  | 2.06 | 0.16 |  |
|  | Control group | Before | 2.12 | 0.04 | |  | 2.12 | 0.04 | |  | 2.12 | 0.05 |  |
|  |  | After | 2.25 | 0.24 | |  | 2.23 | 0.24 | |  | 2.24 | 0.23 |  |
